# Radiation damage to a three-dimensional hydrogel model of the brain perivascular niche

**DOI:** 10.1101/2025.02.20.639287

**Authors:** Y.I. Ivanova, A.C. Nunes, V. Cruz, K. Selting, B.A.C. Harley

**Author notes:** This manuscript conforms with all ethics and integrity policies of the journal. **Corresponding Author:** B.A.C. Harley, Dept. of Chemical and Biomolecular Engineering, Cancer Center at Illinois, Carl R. Woese Institute for Genomic Biology, University of Illinois at Urbana-Champaign, 110 Roger Adams Laboratory, 600 S. Mathews Ave, Urbana, IL 61801.

## Abstract

Glioblastoma (GBM) is a highly aggressive and recurrent brain cancer characterized by diffuse metastasis at the tumor margins. Radiation therapy is a standard component of current treatment and offers potential for improved patient outcomes. While radiation therapy targets GBM cells in the tumor margins, it may also significantly damage adjacent non-cancerous tissues, leading to reduced quality of life and potentially creating a tumor-supportive microenvironment. The perivascular niche (PVN) in the tumor margins is believed to play a significant role in regulating the glioblastoma stem cell subpopulation as well as serving as a site for cancer recurrence and migration. Understanding the impact of radiation on the PVN can better inform radiation schemes and improve our understanding of GBM recurrence, but is difficult *in vivo*. Here we adapt a previously developed three-dimensional hydrogel model of the brain perivascular niche to investigate the impact of radiation dosage and delivery rate on perivascular niche properties *in vitro*. Effects of radiation on vessel architecture can be measured in this hydrogel-based model, suggesting an approach that can provide insight into the effects of radiation on a shorter time scale relative to *in vivo* experiments.

**IMPACT STATEMENT:** Glioblastoma (GBM) is a highly aggressive and recurrent brain cancer characterized by diffuse metastasis at the tumor margins. The perivascular niche (PVN) in the tumor margins plays a significant role in GBM progression and is a target for radiation therapy. We report a method to use three-dimensional hydrogel models of the brain perivascular niche to benchmark the impact of radiation dosage and delivery rate on perivascular niche properties *in vitro*. This approach provides new insight into the effects of radiation on a shorter time scale relative to *in vivo* experiments.

## 1. Introduction

Glioblastoma (GBM) is the most common primary malignant brain tumor with a median survival time of approximately one year and a five-year survival rate of only 5.5% ^1-3^. The current standard of care, known as the Stupp regimen, involves maximal surgical resection, chemoradiotherapy, and adjuvant oral chemotherapy, which has demonstrated improved GBM prognosis since its introduction in 2005 ^4,5^. Despite these advancements in treatment, managing GBM remains difficult due to its aggressive, diffuse, and recurrent nature ^2,3,5,6^. New therapeutic strategies, including the use of novel small drugs, immunotherapy motifs, or modifying the Stupp regimen, are essential to extending median survival time past 15 months ^2,7^. Of particular interest are alterations to radiation protocols to boost efficacy against recurrent tumors and alter the immune response to tumor cells following radiation ^7-12^. With the advent of highly conformal radiation therapy machines, allowing submillimeter precision, the ability to safely alter the dose per fraction within a treatment protocol and avoid permanent, life-altering late effects of radiation has dramatically expanded in recent years. However, this change also introduces an increased risk of a geographic miss leading to tumor recurrence, especially relevant for highly invasive tumors such as GBM.

One of the major challenges in radiation therapy lies in determining appropriate dose per fraction and total dose. While higher total radiation doses can improve efficacy against most cancerous cells, excessive doses come at a steep cost, notably brain necrosis ^6,13^. Necrosis is of greater concern with high dose per fraction (hypofractionated) protocols which can be more damaging to late-responding tissues such as the brain. Thus, it is vital to optimize the radiation dose to achieve maximal therapeutic benefits while minimizing adverse effects. Different radiation delivery methods, such as hypo and hyperfractionation, have shown varying efficacy in GBM treatment ^8,14,15^. Beyond improved targeting of GBM cells, developing effective treatment strategies involves improving our collective understanding of the impact of radiation on the tumor microenvironment. Given resection of the primary tumor mass and subsequent radiation targeting of the tumor margins is part of standard treatment, improved understanding of the role of radiation on the perivascular niches (PVNs) that exist at GBM tumor margins is valuable. The PVN serves as a crucial signaling hub for tumor growth, treatment resistance, and phenotypic changes ^9,16-19^. Notably, a subset of glioblastoma stem cells (GSCs) is believed to play a significant role in treatment resistance and tumor recurrence ^20,21^. The environment of tumor margins, particularly the PVN, is thought to strongly regulate GSC survival, invasion, and therapeutic resistance ^22,23^.

While recent advances in spatial omics and digital pathology are beginning to provide tools to study of the effects of radiation on the brain microenvironment *in vivo* ^24^, there is also an opportunity for experimental platforms that offer physiological relevance as well as reproducible control over experimental conditions. Traditional 2D cell cultures and mouse models have limitations in replicating the PVN’s intricate interactions ^25-29^. Advances in microphysiological models offer an opportunity to create 3D models of the GBM tumor environment via the combination of PVN-associated cells with key components of the brain extracellular matrix. These include efforts to examine the role of the tumor immune microenvironment and chemoradiotherapy on GBM cells in 3D hydrogels ^15,18,30,31^. Recently, we described a 3D hydrogel model of the GBM PVN, combining brain-specific microvascular endothelial cells (ECs), pericytes (PCs), and astrocytes (ACs) with GBM cells to investigate the process of GBM invasion and drug response ^32-34^. We reported successful co-culture of human brain microvascular ECs, human brain vascular PCs, and normal human ACs in a methacrylamide-functionalized gelatin (GelMA) hydrogel to form endothelial networks that mimicked features of the PVN in the brain. We also established a visualization and analysis pipeline to quantify features of the resultant networks such as branch vs. junction parameters ^32,35,36^. Here, we present a methodology for studying radiation effects on the model PVN. We evaluate the impact of different ionizing radiation dose and delivery rates on endothelial network formation and tumor growth. Models that can help resolve the effect of radiation dose on the tumor microenvironment and PVN will provide important, complementary information for the development of more effective radiotherapy strategies for GBM patients.

## 2. Methods

### 2.1. Methacrylamide-Functionalized Gelatin (GelMA) Synthesis

GelMA macromer fabrication used previously published methods ^32,37,38^. Briefly, porcine gelatin type A, 300 bloom (Sigma Aldrich) was dissolved in carbonate buffer at 50°C. 40 µL methacrylic anhydride (Sigma Aldrich) was added dropwise per gram of gelatin. The reaction proceeded for 2 hours with vigorous stirring (500 RPM) and was quenched with 40 mL of 50°C DI water per gram of gelatin. The pH of the solution was adjusted to be between pH of 6-7 through dropwise addition of HCl (Sigma Adlrich). The mixture was then transferred to 12-14 kDA dialysis membranes (Themo Fisher) and dialyzed for seven days against deionized water with daily water changes at 50 °C. The material was then frozen at −20°C and lyophilized. 1HNMR was used to determine the degree of functionalization (DOF). GelMA with DOF between 70% and 80% was used in this study.

### 2.2. Cell Culture

#### 2.2.1. 2D Cell Culture

Human brain microvascular endothelial cells (Cell Systems) were cultured on Attachment Factor-coated (Cell Systems) flasks in Endothelial Growth Medium 2 (Lonza) supplemented with plasmocin prophylactic. Human brain vascular pericytes and normal human astrocytes (Sciencell), were cultured on poly-L-lysine (Sigma Aldrich) coated flasks in Pericyte Growth Medium and Astrocyte Growth Medium (Sciencell) respectively. Cells were expanded for one passage when received from the manufacturer then frozen down. For experiments, the cells were grown until 80% confluency in T-75 flasks (Thermo Fisher) and then seeded for experiments. Cells were lifted using TrypLE (Thermo Fisher). Astrocytes, pericytes, and endothelial cells were used at passages 3, 3, and 5 respectively for all experiments. U87-MG cells were cultured in DMEM with 10% FBS and 1% penicillin/streptomycin. Cells were cultured at 37 °C and 5% CO_2_ per previously published methods ^32^.

#### 2.2.2. Vascular Cell Culture in GelMA Hydrogels

5 wt% GelMA was dissolved in sterile PBS (Thermo Fisher) at 65°C. Lithium acylphosphinate (0.1% w/v, Sigma Aldrich) was added as a photoinitiator ^32,39^. A 3:1:1 Endothelial:Pericyte:Astrocyte cell ratio was used with 1×10^6^ endothelial cells/mL ^32^. Polymer solution and cells were mixed thoroughly and pipetted into circular Teflon molds (5 mm diameter, 1 mm thick. 20 µL volume) and polymerized under UV light (λ = 365 nm, 5.79 mW/cm2, 35 s or λ = 365 nm, 7.27 mW/cm2, 30 s). Hydrogels were cultured in Endothelial Growth Medium 2 supplemented with 50 ng/mL VEGF (Peprotech) for up to twelve days with daily media changes.

#### 2.2.3. GBM-Vascular Co-Culture in GelMA Hydrogels

A 3:1:1:1 Endothelial: Pericyte: Astrocyte: GBM cell ratio was used with 1 × 10^6^ endothelial cells/mL ^32^. Cells were thoroughly mixed in the pre-polymer solution and polymerized under UV light (λ = 365 nm, 7.27 mW/cm2, 30 s). Hydrogels were cultured for up to twelve days in 750 µL of Endothelial Growth Medium 2 + 50 ng/mL VEGF media. When conditioned media collection was desired, 500 µL of the media was collected from each sample before media replacement. Media was collected every 24 hours during the daily media change until the experimental endpoint was reached. Conditioned media was sterile filtered and stored at - 80 °C.

### 2.4. Radiation Exposure

Vascular cell culture hydrogels were irradiated using a Varian™ TrueBeam™ linear accelerator (Varian Medical Systems, Palo Alto, CA, USA) 24 hours after hydrogel formation. Photon beam energy was 6MV and plates containing hydrogels were positioned on 1.5 cm of solid water (Product numbers 805584 and 805588, Gammex Inc, Middleton, WI, USA) to create adequate dose build up with the radiation delivered from beneath the samples. GBM-Vascular Co-Culture hydrogels were exposed to radiation 5 days after hydrogel formation as introduction of GBM cells disrupted network formation ^34^. Radiation doses varied between 0 Gy (a sham control simply transported to and from the facility) and up to 10 Gy. Standard dose delivery rate was set to 6 Gy (600 cGy) per minute while low dose delivery rate was set to 40 cGy/min.

### 2.5. Immunofluorescent Staining

Hydrogels were fixed using formalin for 10 minutes, permeabilized using 0.5% Tween 20 for 10 minutes at room temperature, and blocked using 5% donkey serum (Sigma Aldrich) diluted in 1x PBS (Thermo Fisher) containing 0.1% Tween 20 for 2 hours ^32,35,36,40^. Primary antibodies diluted in the 5% donkey serum solution were then added. Primary antibodies included CD31 (1:100 R&D Systems); PDGFRβ (1:40, R&D Systems) and GFAP (1:500, Proteintech). Donkey Alexa Fluor 488, 568, and 647 were used as secondary antibodies (1:500, Thermo Fisher Scientific). All antibodies were applied overnight at 4 °C, and cells and hydrogels were washed with PBS + 0.1% Tween 20 four times for twenty minutes at room temperature on a rocker between antibody incubations. Hoescht (1:1000, Thermo Fisher Scientific) was used as a nuclear marker.

### 2.6. Vessel Imaging and Metric Analysis

Using 10x magnification, z-stacks with an overall thickness of 200 µm and step size of 5 µm were obtained using a DMi8 (Leica Systems) to extract vessel metrics ^32,35,36,40^. All settings were kept consistent from image to image within an experiment. Images were binarized using the TubeAnalyst Macro referenced in Ngo et al.^32^. Skeletons were then analyzed for metrics including total vessel length, average branch length, number of branches, and number of junctions using a Matlab code described by Crosby et al.^41^. Total vessel length, number of branches, and number of junctions were normalized to unit volume.

### 2.7. Statistics

Statistics were performed using GraphPad Prism (version 10.0.0). All graphs were also made in GraphPad Prism. ROUT outlier test was performed on all data sets with a Q value set to 0.1% to ensure a maximally strict threshold for defining outliers. Gaussian distribution was not assumed for any test. Comparisons between two unpaired groups were performed using a Mann-Whitney U-test, while comparisons between multiple groups were performed using a two-way ANOVA when assumptions were met with radiation and time being used as the two factors. Tukey’s post-hoc test was used to compare significance between these groups. Significance was determined as p < 0.05. All quantitative analyses were performed on hydrogels set up across at least three biological replicates.

## 3. Experiment

### 3.1. Visualization and Analysis of Endothelial Networks in the Vascular Hydrogel Model Are Affected by Hydrogel Permeabilization Time

Perivascular networks were visualized within GelMA hydrogels via fluorescent staining for ECs, PCs, and ACs. Endothelial network architecture parameters (e.g., total network length, average branch length, number of branch points, and number of vessels) were defined via a custom MATLAB code from skeletonized vessel networks created using the EC fluorescent signal (**Figure 1**) ^41^. We first examined the effect of permeabilization time and temperature, blocking solution concentration and incubation time, primary antibody stain (dilutions, duration, wash), secondary antibody stain (duration), and confocal settings on achieving robust metrics of endothelial cell networks. Permeabilization time was found to most strongly affect endothelial network quantification (**Figure 2**). Examining the maximum intensity projections of the binary skeletons generated by our ImageJ pipeline helped understand the effect of permeabilization time. While appearing more complex, the 15 min permeabilization condition also contained a high presence of image artifacts (**Figure 2E**) that were significantly reduced with a shorter (10 min) permeabilization time. As a result, subsequent analyses were performed using a 10 min permeabilization time to ensure reliable vascular metrics, with all other elements of the analysis pipeline kept consistent with previous works ^32^. As a result, network quantification is best used as a relative measurement between experiments as changes in staining conditions can alter the metrics.

**Figure 1.**
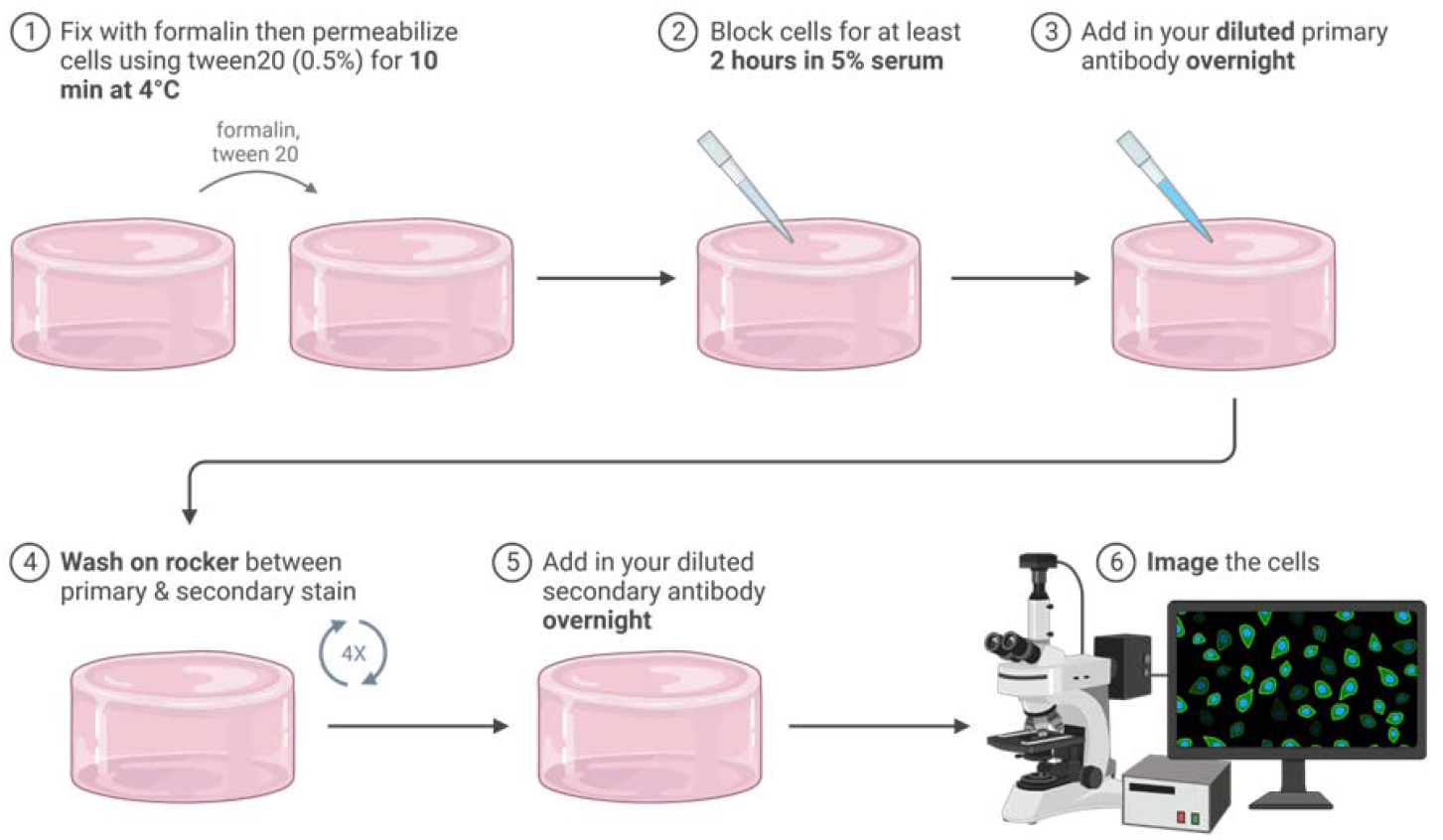
Procedure for staining vascular hydrogels to visualize endothelial networks through antibody staining. The outlined steps represent a summary of the optimized staining protocol. Anything in bold represents a part of the protocol that was tested or optimized. This diagram was made in BioRender.

**Figure 2.**
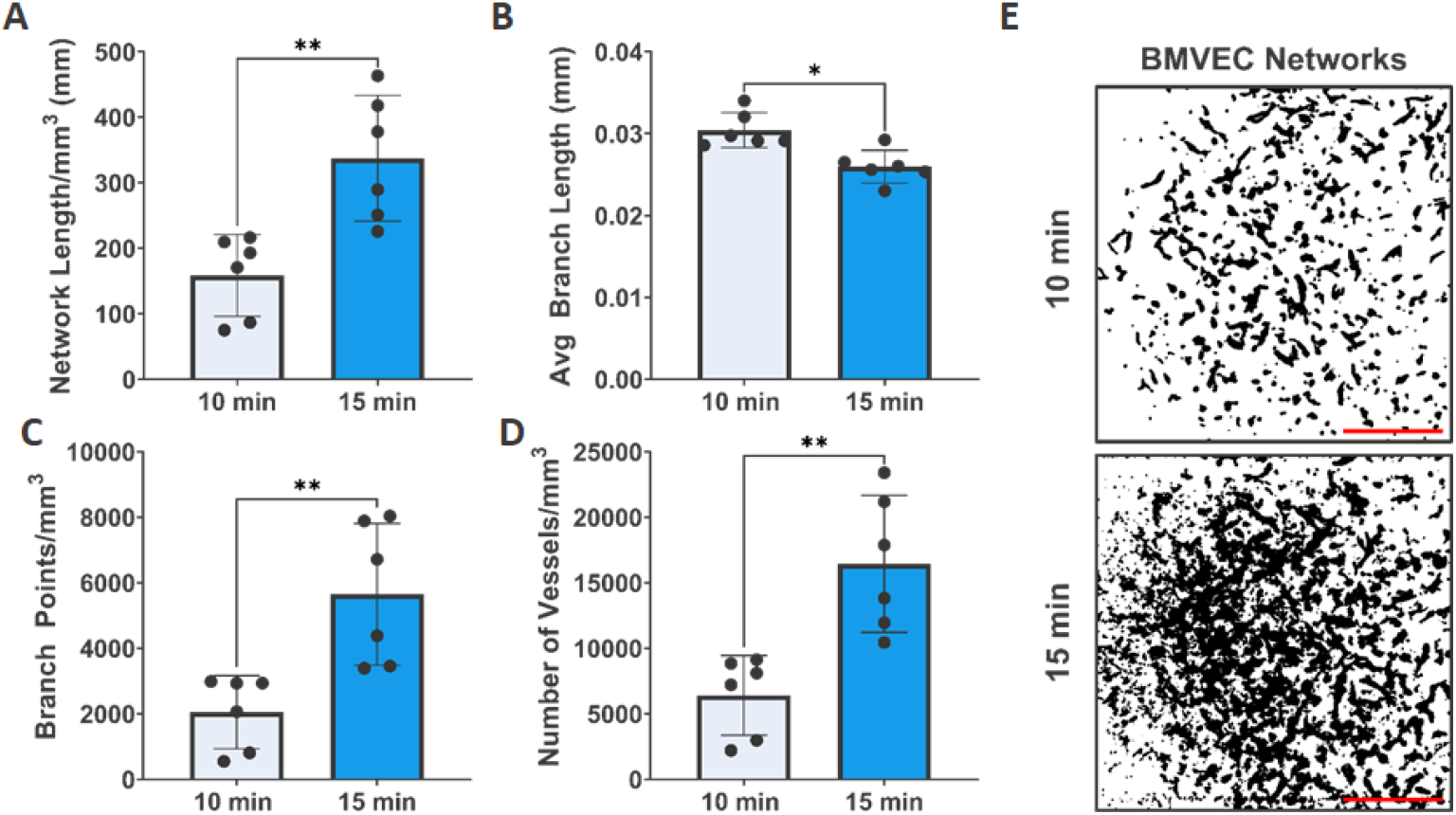
Changing the permeabilization time significantly impacted vessel quantification metrics. **A.** Network length was measured to be significantly longer in the 15 minute group. **B**. Average branch length was measured to be significantly shorter in the 15 minute permeabilization group. **C**. Branch points were measured to be significantly longer in the 15 minute group. **D**. Number of Vessels were measured to be significantly higher in the 15 minute group. **E**. Representative images of the maximum intensity projections of the skeleton z stacks. Error bars represent standard deviation. Significance was determined using a Mann-Whitney U test. * p ≤ 0.05; ** p ≤ 0.01. Scale bar represents 500 µm. Each point represents the average of 6 different locations or images for a single hydrogel (n = 6 total hydrogels).

### 3.2. Improving Endothelial Network Formation in the Vascular Hydrogel Model by Altering Polymer Crosslinking and Density

We subsequently defined the role of the hydrogel microenvironment on the reproducibility of the resultant brain microvascular endothelial networks. All combinations of experimental variables resulted in stable 3D hydrogels with network elastic properties in the range of 2-5 kPa across multiple batches of photoinitiator (PI; **Figure S1**), and were consistent with prior gelatin hydrogels used to replicate the GBM tumor microenvironment ^32^. We examined the effect of hydrogel density (4 vs. 5 wt% GelMA) as well as UV photopolymerization intensity (5.79 vs. 7.27 mW/cm^2^) ^22,32^, using a UV wavelength of 365 nm, outside the wavelength range where it would cause significant damage to cells ^42,43^. We observed significant effects of hydrogel density (4 vs. 5 wt%) on some metrics of endothelial network complexity, notably significantly reduced number of vessels and branch points and significantly increased length of each branch, but no significant difference in overall endothelial network length, a primary indicator of endothelial network presence (**Figure 3A**). However, 4 wt% GelMA hydrogels crosslinked at a UV intensity of 7.27 mW/cm^2^ became more difficult to handle and manipulate with extended culture. As a result, we then examined the effect of UV intensity (5.79 mW/cm^2^ for 30 sec. vs. 7.27 mW/cm^2^ for 30 sec) on endothelial network parameters in 5 wt% GelMA hydrogels. Here, we observed a significant effect of UV exposure on endothelial network parameters (**Figure 3B**), with more robust networks formed in response to lower UV crosslinking intensity after 7 days of culture. These results suggest that UV crosslinking intensity has a stronger influence on perivascular network vessel formation than polymer weight percent of our GelMA hydrogels.

**Figure 3.**
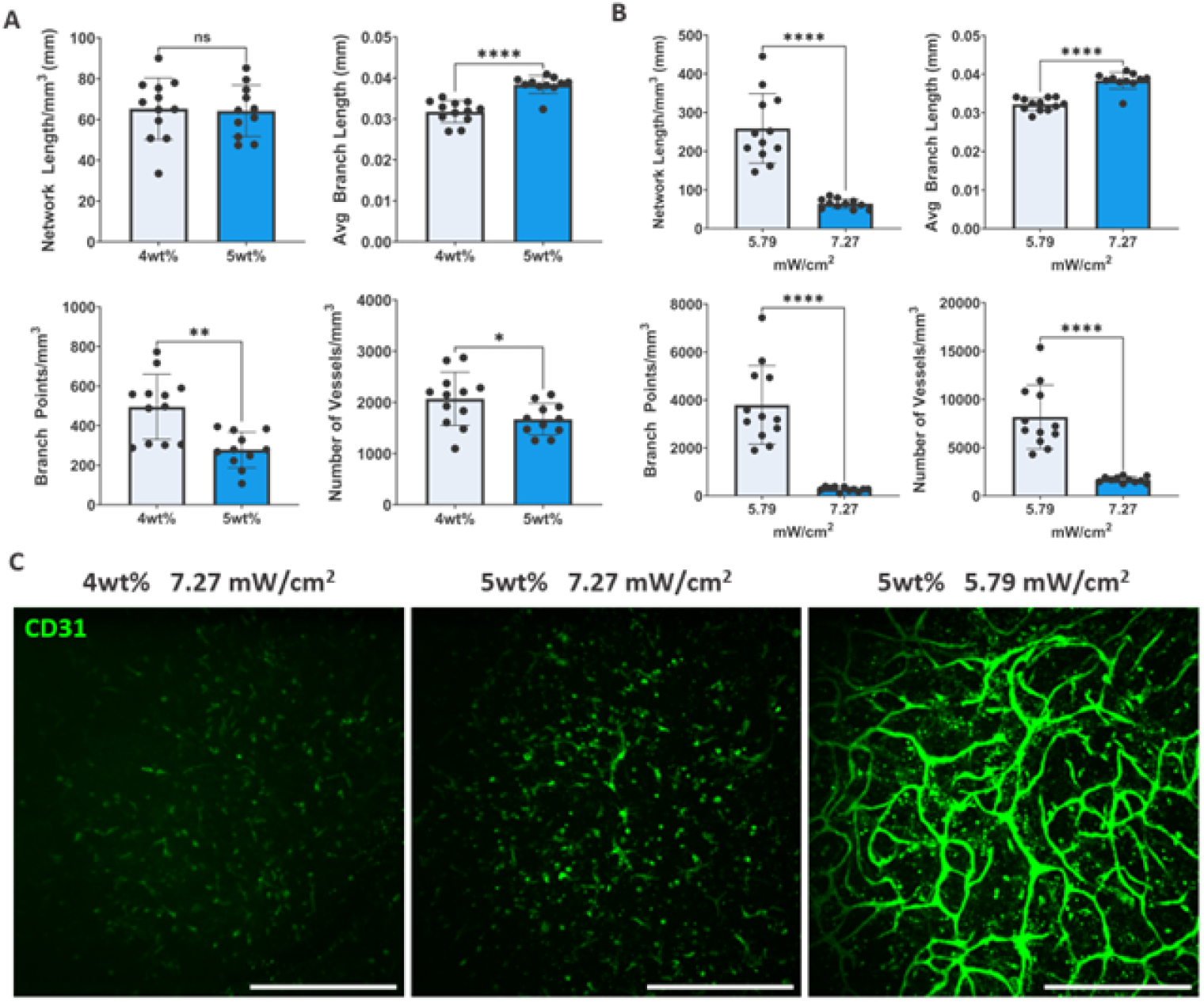
Vessel formation improves with decreased UV intensity. **A.** Network length was the only metric that did not change with alterations in wt%. Average branch length was slightly longer for the 5wt% group. The number of branch points was lower for the 5wt% group. The number of individual vessels was also lower for the 5 wt% group. **B**. Network length metrics changed drastically when UV intensity was altered. Network length decreased significantly for the 7.27 mW group. Average branch length was longer for the 7.27 mW group. The number of branch points was significantly lower for the 7.27 mW group. The number of individual vessels was also lower for the 7.27 mW group. **C**. Representative images shown. Error bars represent standard deviation. All images are shown 7 days post vessel formation. Significance was determined using a Mann-Whitney U test. NS p > 0.05; * p ≤ 0.05; ** p ≤ 0.01; **** p ≤ 0.0001. Each point represents one of 6 technical replicates per hydrogel. N = 3 for total number of hydrogels. Images are maximum intensity projections of 200 µm z stacks taken 5 µm apart. Scale bar represents 500 µm.

### 3.3. Changes in Endothelial Networks Due to Radiation Are Observed at Later Time Points

We then quantified the impact of radiation dose and delivery across 4 different radiation conditions. ECs, PCs, and ACs were seeded in 5 wt% hydrogels and polymerized at 5.79 mW/cm^2^ for 35 seconds. Hydrogels were allowed to culture for one day before exposure to 0 Gy, 5 Gy, or 10 Gy of radiation with the latter two groups being delivered at both a low and standard dose rate (40 vs. 600 cGy/min). Specimens were fixed at 4 or 7 days after radiation exposure (**Figure 4A-D**). A control group was also fixed 1 day after radiation exposure. More significant differences were observed between groups 7 days after radiation exposure, consistent with the time frame for endothelial network formation *in vitro* ^34^. Generally, the 0 Gy group had more robust networks than other groups at day 7, with more variable results at day 4. By day 7, significant effects of radiation were seen on the resultant endothelial cell networks, with some trends associated with radiation dose and rate. The control condition (0 Gy) resulted in the most robust networks, with evidence of budding networks in both the 5 Gy low and 10 Gy low dose rate conditions when compared to their respective standard dose rate conditions (**Figure 4E**). In general, exposure to radiation resulted in significantly reduced total endothelial network lengths, reduced numbers of branch points in the networks, and reduced numbers of vessels. There were no apparent changes in endothelial network parameters between normal and low exposure rates at 5 Gy dose. And while not often significant, low dose rate radiation showed less of an effect on endothelial network parameters (more similar to 0 Gy control) for the 10 Gy dose groups. The lack of more statistical comparisons is likely due to the large standard deviations seen in irradiated endothelial networks.

**Figure 4.**
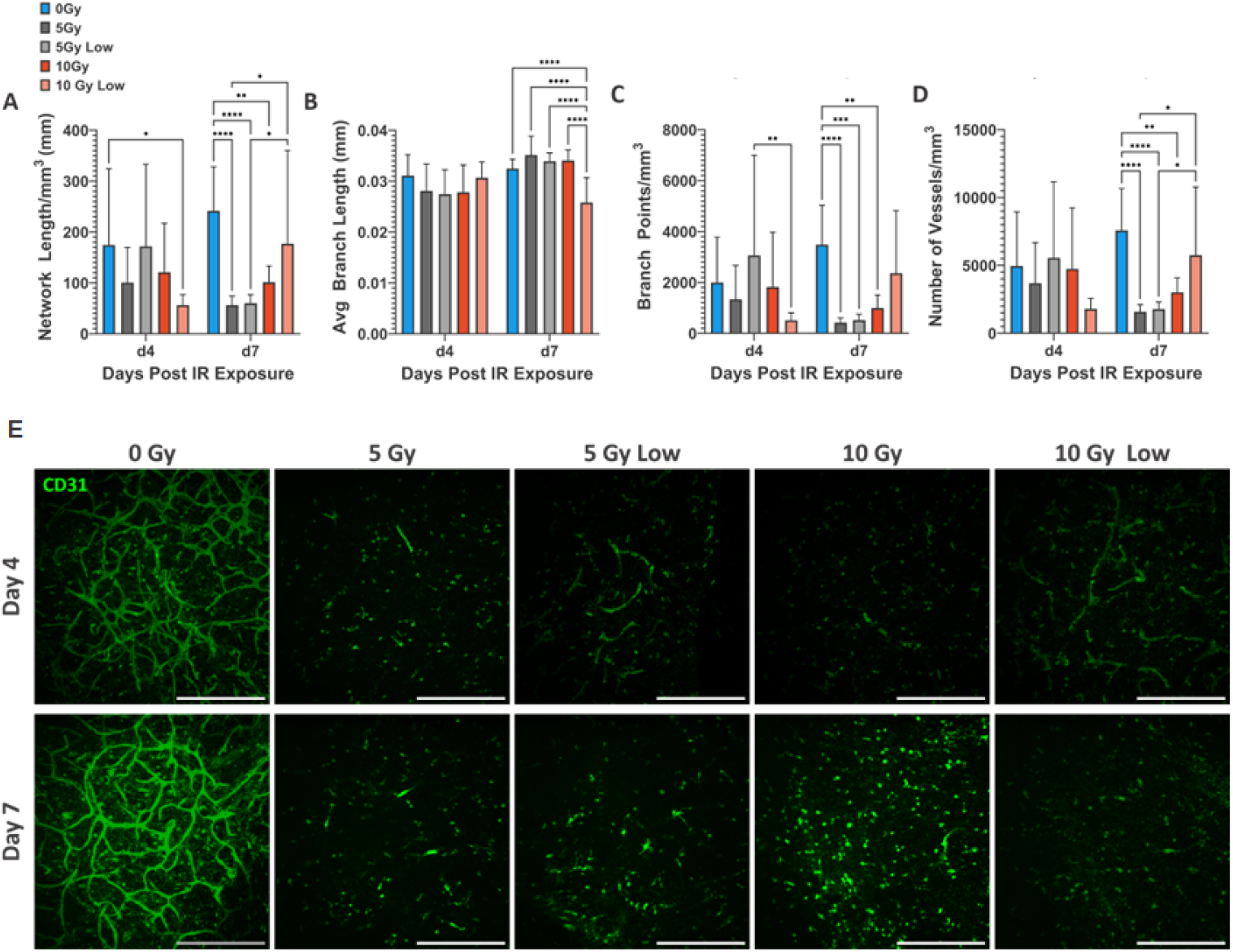
Vascular metrics had more significant differences between groups 7 days post IR exposure: 0Gy vs 10Gy at 40 (*low*) vs. 600 (*standard*) cGy/min. **A.** 0Gy vasculature network length was significantly longer than the 10Gy low group at day 4. 0Gy network length was also significantly longer than all groups except 10Gy low at day 7. **B**. Average branch length of 10Gy low group was significantly shorter at day 7. **C**. 0Gy branch points were significantly higher than all groups except the 10Gy low group at day 7. **D**. 0Gy network length was significantly longer than all groups except 10Gy low at day 7. 10Gy low group was significantly higher than both the 5Gy and 5Gy low group. **E**. Representative images shown. Significance was determined using a Tukey’s post-hoc test after a two-way ANOVA. Only significance between groups at the same time point are shown to avoid interaction effects between time and radiation. * p ≤ 0.05; ** p ≤ 0.01; *** p ≤ 0.001; **** p ≤ 0.0001. N = 3 total hydrogels with 6 distinct images per gel. Images are maximum intensity projections of 200 µm z stacks taken 5 µm apart. Scale bar represents 500 µm.

### 3.4. Including GBM cells in irradiated endothelial cell networks

We subsequently examined the effect of radiation on endothelial networks in the presence of GBM cells. ECs, PCs, ACs, and U87-MG GBM cells were seeded in 5 wt% GelMA hydrogels (7.27 mW/cm^2^ for 30 seconds), chosen to mimic a stiffer brain tumor microenvironment ^44-47^. Hydrogels were cultured for five days before being exposed to either 0 Gy or 10 Gy of radiation at the standard dose rate (600 cGy/min); endothelial networks were subsequently quantified at 1, 4, and 7 days after radiation exposure (6, 9, and 12 days after initial hydrogel formation). Significant differences were only observed 1 day after irradiation, with increased numbers of branches, vessels, and vessel length in irradiated specimens. Additionally, vessel networks in irradiated specimens showed a negative trend with time, with no significant reductions in numbers of branches, vessels, and overall network length. In comparison, non-irradiated (0 Gy) samples generally showed upward trends in these variables with time, suggesting more mature vessels (**Figure 5B-E**).

**Figure 5.**
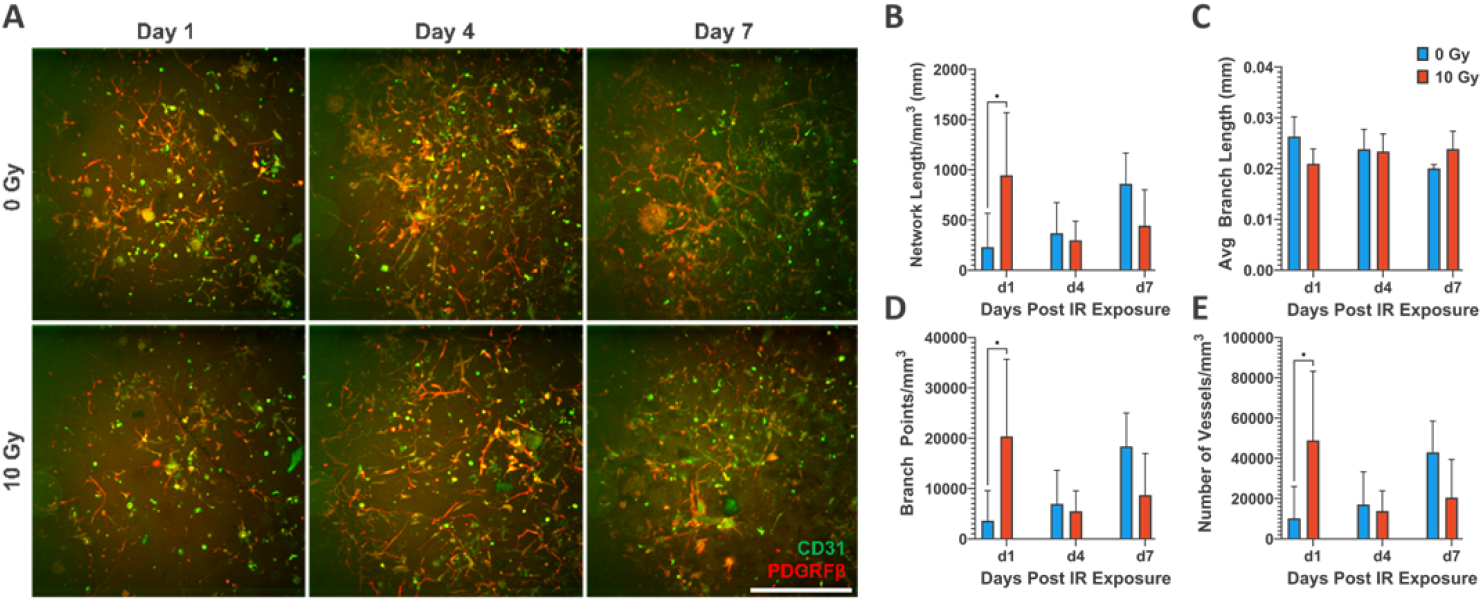
Inclusion of disperse GBM cells may obscure network differences. **A.** Representative of images of vascular networks at different time points under different radiation conditions. **B**. There is a significant difference in network length between the 0 Gy and 10 Gy group on day 1. **C**. There are no differences between average branch length for any groups at any time point. **D**. There is a significant difference in branch points between the 0 Gy and 10 Gy group on day 1. **E**. There is a significant difference in network length between the 0 Gy and 10 Gy group on day 1. CD31 is a marker for endothelial cells. PDGRFβ is a marker for pericytes. Images are maximum intensity projections of 200 µm z stacks taken 5 µm apart. Scale bar represents 500 µm. Error bars represent standard deviation. Significance was determined using a Mann-Whitney U test. * p ≤ 0.05. N = 3 total hydrogels with 6 images per gel.

## 4. Discussion

An improved understanding of radiation damage to the margins of the glioblastoma tumor microenvironment, particularly to the PVN, is essential to better understand glioblastoma recurrence and drug sensitivity. Here we report a method to adapt a three-dimensional hydrogel to evaluate the effect of radiation exposure on metrics of PVN maturation in the presence or absence of GBM cells. This methods-centered report described modifications to the fabrication scheme of a 3D perivascular hydrogel model to provide reproducible vessel models as an essential precursor to studies that evaluate the effect of radiation dose on a shorter time scale than possible in traditional *in vivo* models. Unlike 2D or Matrigel-based models, this approach incorporates the structural complexity of the PVN while providing precise control over variables such as cell type, stiffness, and extracellular matrix composition. Given that this perivascular hydrogel model has previously been shown to evaluate the effects of multi-cell signaling on glioblastoma cell invasion, retention of a stem like phenotype, and metrics of GBM cell motility ^32,48^, the ability to now examine the effect of radiation exposure on the engineered PVN suggests important new avenues for future study.

A major advantage of this approach is that microphysiological hydrogel models provide an avenue for rapid and robust investigations of PVN network complexity and GBM cell response over short-time scales (days to weeks) ^49^. And as opposed to prior efforts using tissue-agnostic cells lines (e.g., human umbilical vein endothelial cells and human lung fibroblasts), here we provide an approach to create networks from brain-specific sources (brain microvascular endothelial cells, pericytes, and astrocytes) in a readily degradable hydrogel model to facilitate rapid cell isolation and analysis, and using a gelatin macromer amenable to extracellular matrix additions ^38,50-54^. Notably, we observe that exposure to radiation (0 – 10 Gy) and changes in dose rate (*low* 40 vs. *standard* 600 cGy/min) alter network response over time. Including the U87-MG cell line within the model may obscure some findings, but offers an exciting avenue for future work. Notably, co-culture of GBM cells with endothelial networks has already been shown to induce processes of co-option and regression characteristic of the native GBM margins ^34^. While the presence of GBM cells can induce vessel regression in our hydrogel model, the addition of radiation could negatively affect the vessel networks directly but may indirectly help the networks by reducing the deleterious effect of co-encapsulated GBM cells. This suggests future experiments to consider temporal patterns of endothelial network complexity in response to these competing stimuli. Such studies are largely intractable *in vivo* due to technical limitations associated with repeatedly and reproducibly monitoring microvascular network morphology changes, further highlighting the opportunity to leverage this endothelial network-GBM model described here in future extensions of this work.

A primary limitation of this work is the need to correlate findings *in vitro* to functional changes *in vivo*. Certainly, changes to perivascular network complexity as well as the stability of tight junctions within these networks will play a significant role in the local tumor microenvironment that may alter the activity of GBM cells post radiation exposure. Future studies should also therefore correlate changes in vascular architecture seen here *in vitro* to shifts in functional phenotype *in vivo*. Indeed, we recently reported an approach to use Fourier transform infrared (FTIR) spectroscopic imaging for label-free detection of biomolecular changes in the brain 15 month following 30-Gy RT ^24^. Such observations provide a route to consider functional changes in the brain vascular microenvironment beyond characterization of vessel morphology that could be complementary to the methodology reported here. Future studies should also consider a wider range of GBM tumor microenvironment associated cells. We have previously shown that microglia play a significant role in GBM cell proliferation and invasion in hydrogel models of the brain tumor microenvironment ^55^. Further, their ability to catalyze a larger neuroinflammatory response is important to understand in the context of radiation exposure. Other limitations of the work include that choice of diffusely seeded GBM and vascular cells; future endeavors could consider the introduction of GBM cells, particularly GBM spheroids ^32^, into the PVN after it forms. Regardless, we report a methodological approach to assemble defined 3D hydrogel models of the brain perivascular environment in order to assess changes in perivascular network parameters in response to clinically relevant radiation exposure. Ongoing and future research directions for the project include inclusion of patient-derived xenograft cell lines into the PVN model to evaluate patterns of drug sensitivity and invasion, inclusion of microglia, as well as efforts to consider the effect of hypoxia.

New clinical strategies to improve patient outcomes for glioblastoma require innovative therapeutic strategies. Particularly optimizing radiation dosages and understanding its impact on the tumor microenvironment can allow an existing treatment to be leveraged more effectively. This study introduced a 3D hydrogel model to evaluate the effect of radiation exposure on metrics of PVN architecture. Significant findings highlight the critical role of hydrogel permeabilization time in reproducibly measuring 3D PVN architecture, the greater influence of UV intensity versus polymer weight in hydrogel endothelial network formation, and the prominence of radiation-induced changes in vascular networks at later (4 vs. 7 days post radiation) time points. These insights serve as a foundation for future investigations aimed at tailoring radiation-based treatments for GBM, potentially enhancing therapeutic efficacy and patient outcomes.

## Acknowledgements

The authors would like to acknowledge the following institutes for access to their facilities and services: the Carl R. Woese Institute for Genomic Biology, the Tumor Engineering and Phenotyping Shared Resource (TEP) at the Cancer Center at Illinois, and the Roy J. Carver Biotechnology Center (Flow Cytometry Facility, UIUC) for assistance with design of flow cytometry panels. Research reported in this publication was supported by the National Cancer Institute under Award Number R01 CA256481 (BACH). We are also grateful for funds provided by the NSF Graduate Research Fellowship (DGE-1746047 to YII). Additional support was provided by the Carl R. Woese Institute for Genomic Biology and the Chemical and Biomolecular Engineering Dept. at the University of Illinois at Urbana-Champaign. The interpretations and conclusions presented are those of the authors and are not necessarily endorsed by the National Institutes of Health or the National Science Foundation.

## Contributions (CRediT: Contributor Roles Taxonomy)

**Y. Ivanova:** Conceptualization, Data curation, Formal Analysis, Visualization, Investigation, Methodology, Writing – original draft, Writing – review & editing. **A. Nunes:** Investigation, Formal Analysis. **V. Cruz:** Investigation, Formal Analysis. **K. Selting:** Conceptualization, Resources, Supervision, Writing – review & editing. **Brendan Harley:** Conceptualization, Resources, Project administration, Funding acquisition, Supervision, Writing – review & editing.

**Figure S1.**
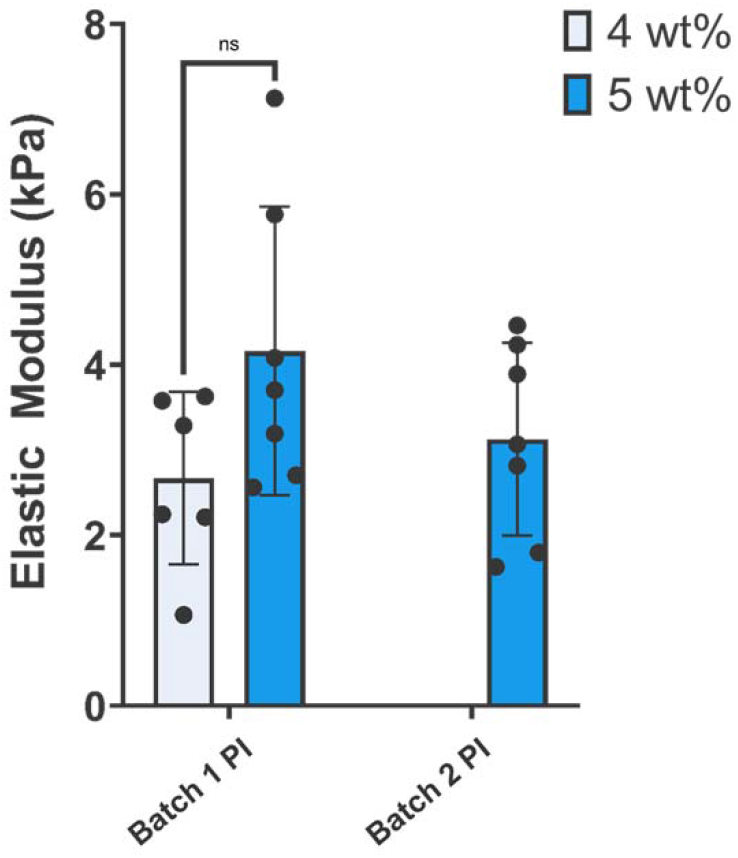
There is no significant difference in stiffness when polymer weight percent is altered. Error bars represent standard deviation. Significance was determined using a Mann-Whitney U test. ns p > 0.999. Each point represents an individual hydrogel (n = 6).

## Notes

### Competing Interest Statement

The authors have declared no competing interest.

## References

1. Wu W, Klockow JL, Zhang M, et al. Glioblastoma multiforme (GBM): An overview of current therapies and mechanisms of resistance. Pharmacol Res 2021;171(105780, doi:10.1016/j.phrs.2021.105780

2. Davis ME. Glioblastoma: Overview of Disease and Treatment. Clin J Oncol Nurs 2016;20(5 Suppl):S2–8, doi:10.1188/16.CJON.S1.2-8

3. Thakkar JP, Dolecek TA, Horbinski C, et al. Epidemiologic and molecular prognostic review of glioblastoma. Cancer Epidemiol Biomarkers Prev 2014;23(10):1985–96, doi:10.1158/1055-9965.EPI-14-0275

4. Stupp R, Mason WP, van den Bent MJ, et al. Radiotherapy plus concomitant and adjuvant temozolomide for glioblastoma. N Engl J Med 2005;352(10):987–96, doi:10.1056/NEJMoa043330

5. Lakomy R, Kazda T, Selingerova I, et al. Real-World Evidence in Glioblastoma: Stupp’s Regimen After a Decade. Front Oncol 2020;10(840, doi:10.3389/fonc.2020.00840

6. Chao ST, Ahluwalia MS, Barnett GH, et al. Challenges with the diagnosis and treatment of cerebral radiation necrosis. Int J Radiat Oncol Biol Phys 2013;87(3):449–57, doi:10.1016/j.ijrobp.2013.05.015

7. Burr AR, Robins HI, Bayliss RA, et al. Outcomes From Whole-Brain Reirradiation Using Pulsed Reduced Dose Rate Radiation Therapy. Adv Radiat Oncol 2020;5(5):834–839, doi:10.1016/j.adro.2020.06.021

8. Minniti G, Niyazi M, Alongi F, et al. Current status and recent advances in reirradiation of glioblastoma. Radiat Oncol 2021;16(1):36, doi:10.1186/s13014-021-01767-9

9. Barker HE, Paget JT, Khan AA, et al. The tumour microenvironment after radiotherapy: mechanisms of resistance and recurrence. Nat Rev Cancer 2015;15(7):409–25, doi:10.1038/nrc3958

10. Ellor SV, Pagano-Young TA, Avgeropoulos NG. Glioblastoma: background, standard treatment paradigms, and supportive care considerations. J Law Med Ethics 2014;42(2):171–82, doi:10.1111/jlme.12133

11. Kleinberg L, Sloan L, Grossman S, et al. Radiotherapy, Lymphopenia, and Host Immune Capacity in Glioblastoma: A Potentially Actionable Toxicity Associated With Reduced Efficacy of Radiotherapy. Neurosurgery 2019;85(4):441–453, doi:10.1093/neuros/nyz198

12. Rivera M, Sukhdeo K, Yu J. Ionizing radiation in glioblastoma initiating cells. Front Oncol 2013;3(74, doi:10.3389/fonc.2013.00074

13. Yang H, Chopp M. Functional Issues in Brain Tumor Treatment. Journal of Neurology & Neurophysiology 2011;01(5), doi:10.4172/2155-9562.S5-002

14. Barani IJ, Larson DA. Radiation Therapy of Glioblastoma. In: Current Understanding and Treatment of Gliomas. (Raizer J, Parsa A. eds.) Springer International Publishing: Cham; 2015; pp. 49–73.

15. Yi HG, Jeong YH, Kim Y, et al. A bioprinted human-glioblastoma-on-a-chip for the identification of patient-specific responses to chemoradiotherapy. Nat Biomed Eng 2019;3(7):509–519, doi:10.1038/s41551-019-0363-x

16. Nowosad A, Marine JC, Karras P. Perivascular niches: critical hubs in cancer evolution. Trends Cancer 2023;9(11):897–910, doi:10.1016/j.trecan.2023.06.010

17. Vilalta M, Rafat M, Giaccia AJ, et al. Recruitment of circulating breast cancer cells is stimulated by radiotherapy. Cell Rep 2014;8(2):402–9, doi:10.1016/j.celrep.2014.06.011

18. Henrik Heiland D, Ravi VM, Behringer SP, et al. Tumor-associated reactive astrocytes aid the evolution of immunosuppressive environment in glioblastoma. Nat Commun 2019;10(1):2541, doi:10.1038/s41467-019-10493-6

19. Krisnawan VE, Stanley JA, Schwarz JK, et al. Tumor Microenvironment as a Regulator of Radiation Therapy: New Insights into Stromal-Mediated Radioresistance. Cancers (Basel) 2020;12(10), doi:10.3390/cancers12102916

20. Alves ALV, Gomes INF, Carloni AC, et al. Role of glioblastoma stem cells in cancer therapeutic resistance: a perspective on antineoplastic agents from natural sources and chemical derivatives. Stem Cell Res Ther 2021;12(1):206, doi:10.1186/s13287-021-02231-x

21. Brooks MD, Sengupta R, Snyder SC, et al. Hitting Them Where They Live: Targeting the Glioblastoma Perivascular Stem Cell Niche. Curr Pathobiol Rep 2013;1(2):101–110, doi:10.1007/s40139-013-0012-0

22. So JS, Kim H, Han KS. Mechanisms of Invasion in Glioblastoma: Extracellular Matrix, Ca(2+) Signaling, and Glutamate. Front Cell Neurosci 2021;15(663092, doi:10.3389/fncel.2021.663092

23. Borovski T, Beke P, van Tellingen O, et al. Therapy-resistant tumor microvascular endothelial cells contribute to treatment failure in glioblastoma multiforme. Oncogene 2013;32(12):1539–48, doi:10.1038/onc.2012.172

24. Chen J, Laverty DJ, Talele S, et al. Aberrant ATM signaling and homology-directed DNA repair as a vulnerability of p53-mutant GBM to AZD1390-mediated radiosensitization. Science translational medicine 2024;16(734):eadj5962, doi:10.1126/scitranslmed.adj5962

25. Fessler E, Borovski T, Medema JP. Endothelial cells induce cancer stem cell features in differentiated glioblastoma cells via bFGF. Mol Cancer 2015;14(157, doi:10.1186/s12943-015-0420-3

26. Berg TJ, Marques C, Pantazopoulou V, et al. The Irradiated Brain Microenvironment Supports Glioma Stemness and Survival via Astrocyte-Derived Transglutaminase 2. Cancer Res 2021;81(8):2101–2115, doi:10.1158/0008-5472.CAN-20-1785

27. Paolillo M, Comincini S, Schinelli S. In Vitro Glioblastoma Models: A Journey into the Third Dimension. Cancers (Basel) 2021;13(10), doi:10.3390/cancers13102449

28. Gomez-Roman N, Chong MY, Chahal SK, et al. Radiation Responses of 2D and 3D Glioblastoma Cells: A Novel, 3D-specific Radioprotective Role of VEGF/Akt Signaling through Functional Activation of NHEJ. Molecular Cancer Therapeutics 2020;19(2):575–589, doi:10.1158/1535-7163.Mct-18-1320

29. Neufeld L, Yeini E, Reisman N, et al. Microengineered perfusable 3D-bioprinted glioblastoma model for in vivo mimicry of tumor microenvironment. Sci Adv 2021;7(34), doi:10.1126/sciadv.abi9119

30. Bylicky MA, Mueller GP, Day RM. Radiation resistance of normal human astrocytes: the role of non-homologous end joining DNA repair activity. J Radiat Res 2019;60(1):37–50, doi:10.1093/jrr/rry084

31. Zhang H, Zhou Y, Cui B, et al. Novel insights into astrocyte-mediated signaling of proliferation, invasion and tumor immune microenvironment in glioblastoma. Biomed Pharmacother 2020;126(110086, doi:10.1016/j.biopha.2020.110086

32. Ngo MT, Sarkaria JN, Harley BAC. Perivascular Stromal Cells Instruct Glioblastoma Invasion, Proliferation, and Therapeutic Response within an Engineered Brain Perivascular Niche Model. Adv Sci (Weinh) 2022;9(31):e2201888, doi:10.1002/advs.202201888

33. Ngo MT, Harley BAC. Perivascular signals alter global gene expression profile of glioblastoma and response to temozolomide in a gelatin hydrogel. Biomaterials 2019;198(122-134, doi:10.1016/j.biomaterials.2018.06.013

34. Ngo MT, Harley BA. The Influence of Hyaluronic Acid and Glioblastoma Cell Coculture on the Formation of Endothelial Cell Networks in Gelatin Hydrogels. Adv Healthc Mater 2017;6(22), doi:10.1002/adhm.201700687

35. Ngo MT, Barnhouse VR, Gilchrist AE, et al. Hydrogels Containing Gradients in Vascular Density Reveal Dose-Dependent Role of Angiocrine Cues on Stem Cell Behavior. Adv Funct Mater 2021;31(51), doi:10.1002/adfm.202101541

36. Zambuto SG, Theriault H, Jain I, et al. Endometrial decidualization status modulates endometrial microvascular complexity and trophoblast outgrowth in gelatin methacryloyl hydrogels. npj Women’s Health 2024;2(1):22, doi:10.1038/s44294-024-00020-4

37. Kriuchkovskaia VA, Eames EK, Riggins RB, et al. Acquired Temozolomide Resistance Instructs Patterns of Glioblastoma Behavior in Gelatin Hydrogels. Advanced Healthcare Materials n/a(n/a):2400779, doi:10.1002/adhm.202400779

38. Pedron S, Hanselman JS, Schroeder MA, et al. Extracellular Hyaluronic Acid Influences the Efficacy of EGFR Tyrosine Kinase Inhibitors in a Biomaterial Model of Glioblastoma. Adv Healthc Mater 2017;6(21), doi:10.1002/adhm.201700529

39. Mahadik BP, Pedron Haba S, Skertich LJ, et al. The use of covalently immobilized stem cell factor to selectively affect hematopoietic stem cell activity within a gelatin hydrogel. Biomaterials 2015;67(297-307, doi:10.1016/j.biomaterials.2015.07.042

40. Barnhouse V, Petrikas N, Crosby C, et al. Perivascular secretome influences hematopoietic stem cell maintenance in a gelatin hydrogel. Ann Biomed Eng 2021;49(2):780–792, doi:10.1007/s10439-020-02602-0

41. Crosby CO, Valliappan D, Shu D, et al. Quantifying the Vasculogenic Potential of Induced Pluripotent Stem Cell-Derived Endothelial Progenitors in Collagen Hydrogels. Tissue Eng Part A 2019;25(9-10):746-758, doi:10.1089/ten.TEA.2018.0274

42. Masuma R, Kashima S, Kurasaki M, et al. Effects of UV wavelength on cell damages caused by UV irradiation in PC12 cells. J Photochem Photobiol B 2013;125(202-8, doi:10.1016/j.jphotobiol.2013.06.003

43. Wong DY, Ranganath T, Kasko AM. Low-Dose, Long-Wave UV Light Does Not Affect Gene Expression of Human Mesenchymal Stem Cells. PLoS One 2015;10(9):e0139307, doi:10.1371/journal.pone.0139307

44. Chen J-W, Pedron S, Harley BAC. The combined influence of hydrogel stiffness and matrix-bound hyaluronic acid content on glioblastoma invasion. Macromol Biosci 2017;17(8):1700018, doi:10.1002/mabi.201700018

45. Miroshnikova YA, Mouw JK, Barnes JM, et al. Tissue mechanics promote IDH1-dependent HIF1[alpha]-tenascin C feedback to regulate glioblastoma aggression. Nat Cell Biol 2016;18(12):1336–1345, doi:10.1038/ncb3429 http://www.nature.com/ncb/journal/v18/n12/abs/ncb3429.html#supplementary-information

46. Heffernan JM, Overstreet DJ, L. LD, et al. Bioengineered scaffolds for 3D analysis of glioblastoma proliferation and invasion. Ann Biomed Eng 2015;43(8):1965–77, doi:10.1007/s10439-014-1223-1

47. Umesh V, Rape AD, Ulrich TA, et al. Microenvironmental stiffness enhances glioma cell proliferation by stimulating epidermal growth factor receptor signaling. PloS one 2014;9(7):e101771, doi:10.1371/journal.pone.0101771

48. Ngo MT, Karvelis E, Harley BAC. Multidimensional hydrogel models reveal endothelial network angiocrine signals increase glioblastoma cell number, invasion, and temozolomide resistance. Integr Biol (Camb) 2020;12(6):139–149, doi:10.1093/intbio/zyaa010

49. Straehla JP, Hajal C, Safford HC, et al. A predictive microfluidic model of human glioblastoma to assess trafficking of blood-brain barrier-penetrant nanoparticles. Proc Natl Acad Sci U S A 2022;119(23):e2118697119, doi:10.1073/pnas.2118697119

50. Neves ER, Anand A, Mueller J, et al. Targeting glioblastoma tumor hyaluronan to enhance therapeutic interventions that regulate metabolic cell properties. bioRxiv 2024, doi:10.1101/2024.01.05.574065

51. Pedron S, Becka E, Harley BA. Regulation of glioma cell phenotype in 3D matrices by hyaluronic acid. Biomaterials 2013;34(30):7408–17, doi:10.1016/j.biomaterials.2013.06.024

52. Pedron S, Becka E, Harley BA. Spatially gradated hydrogel platform as a 3D engineered tumor microenvironment. Adv Mater 2015;27(9):1567–72, doi:10.1002/adma.201404896

53. Pedron S, Polishetty H, Pritchard AM, et al. Spatially graded hydrogels for preclinical testing of glioblastoma anticancer therapeutics. MRS Commun 2017;7(3):442–449, doi:10.1557/mrc.2017.85

54. Pedron S, Wolter GL, Chen JE, et al. Hyaluronic acid-functionalized gelatin hydrogels reveal extracellular matrix signals temper the efficacy of erlotinib against patient-derived glioblastoma specimens. Biomaterials 2019;219(119371, doi:10.1016/j.biomaterials.2019.119371

55. Chen JE, Lumibao J, Leary S, et al. Crosstalk between microglia and patient-derived glioblastoma cells inhibit invasion in a three-dimensional gelatin hydrogel model. J Neuroinflammation 2020;17(1):346, doi:10.1186/s12974-020-02026-6

